# *SWELL1* is a glucose sensor required for β-cell excitability and insulin secretion

**DOI:** 10.1101/155093

**Authors:** Chen Kang, Susheel K. Gunasekar, Anil Mishra, Litao Xie, Yanhui Zhang, Saachi Pai, Yiwen Gao, Andrew W. Norris, Samuel B. Stephens, Rajan Sah

**Affiliations:** Department of Internal Medicine, Division of Cardiovascular Medicine, University of Iowa, Carver College of Medicine, Iowa City, IA, 52242; Department of Pediatrics, University of Iowa, Carver College of Medicine, Iowa City, IA, 52242; Fraternal Order of the Eagles Diabetes Research Center, Iowa City, IA, 52242; Abboud Cardiovascular Research Center, University of Iowa, Carver College of Medicine, Iowa City, IA, 52242

## Abstract

Insulin secretion from the pancreatic β-cell initiated by activation of voltage-gated Ca^2+^ channels (VGCC) to trigger Ca^2+^-mediated insulin vesicle fusion with the β-cell plasma membrane. The firing of VGCC depends on the β-cell membrane potential, which is in turn mediated by the balance of depolarizing (excitatory) and hyperpolarizing (inhibitory) ionic currents^1-3^. While much attention has focused on inhibitory potassium currents^4-10^ there is little knowledge about the excitatory currents required to depolarize the β-cell, including the molecular identity of these excitatory currents^3^. Here we show that SWELL1 (LRRC8a) mediates a swell-activated, depolarizing chloride current (I_Cl,SWELL)_ in β-cells. Hypotonic and glucose-stimulated β-cell swelling activates SWELL1-mediated I_Cl,SWELL_ and this is required for both glucose-stimulated and hypotonic swell-mediated activation of VGCC-dependent intracellular calcium signaling in β-cells. SWELL1 KO MIN6 cells and β-cell targeted SWELL1 KO murine islets exhibit significantly impaired glucose-stimulated insulin secretion, with preserved insulin content *in vitro*. Tamoxifen-inducible β-cell targeted *SWELL1* KO mice have normal fasting insulin levels but display markedly impaired glucose-stimulated insulin secretion. Our results reveal a physiological role for SWELL1 as a glucose sensor - linking glucose-mediated β-cell swelling to SWELL1-dependent activation of VGCC-triggered calcium signaling, and highlights SWELL1-mediated “swell-secretion” coupling as required for glucose-stimulated insulin secretion.

---

Recent studies have identified SWELL1 (LRRC8a) as a required component of a swell-activated Cl^−^ current I_Cl,SWELL_ or ***V***olume-***R***egulated ***A***nion ***C***urrent (VRAC) in common cell lines^11^, ^12^, forming multimeric channels with LRRC8b-e^13^. To determine if SWELL1 is also required for VRAC in pancreatic β-cells we suppressed SWELL1 in mouse insulinoma (MIN6) cells by adenoviral transduction with an shRNA-directed against SWELL1 (Ad-U6-shSWELL1-mCherry; Fig. 1a) as compared to a scrambled shRNA control (Ad-U6-shSCR-mCherry). We observed robust knock-down of SWELL1 protein (Fig. 1b) and a significant reduction in hypotonic swell-activated VRAC in Ad-shSWELL1 relative to Ad-shSCR transduced MIN6 cells (Fig. 1c&d). To determine whether SWELL1 is also required for VRAC in mouse primary β-cells we isolated islets from *SWELL1* floxed mice (*SWELL1^fl^*)^14^ and transduced them with an adenovirus expressing GFP under control of a rat insulin promoter (Ad-RIP2-GFP) to allow positive identification of β-cells (GFP+ cells). *SWELL1^fl^* islets were further treated with adenovirus expressing Cre-mCherry to allow Cre-mediated excision of the floxed SWELL1 allele or control virus expressing mCherry alone (Supplementary Fig. 1a). By selecting GFP+/mCherry+ cells, we are able to patch-clamp either control WT β-cells (*SWELL1^fl^* β-cells) or *SWELL1* KO β-cells (*SWELL1^fl^*/Cre β-cells; Fig. 1e). We find that WT β-cells express substantial swell-activated current and this is entirely abolished upon Cre-mediated recombination in *SWELL1* KO β-cells (Fig. 1f-h). We next tested whether SWELL1 is also required for VRAC in human β-cells and applied a similar approach. We transduced human islets with Ad-RIP2-GFP and Ad-shSWELL1-mCherry or Ad-sh-SCR-mCherry (Supplementary Fig. 1b), in order to isolate and patch-clamp human β-cells (GFP+) subjected to shRNA-mediated SWELL1 KD or to a scrambled control (GFP+/mCherry+; Fig. 1i). Similar to mouse β-cell recordings we find that human β-cells also express significant SWELL1-mediated swell-activated current (Fig. 1j-l). Indeed, in all β-cells patch-clamped, the reversal potential was ~-12 mV, which is near the reversal potential for Cl^−^ under our recording conditions, and thus consistent with SWELL1-mediating a swell-activated Cl^−^ conductance in β-cells^15^^−^^17^. These data demonstrate that SWELL1 is required for VRAC or ICl,SWELL in pancreatic β-cells.

**Figure 1.**
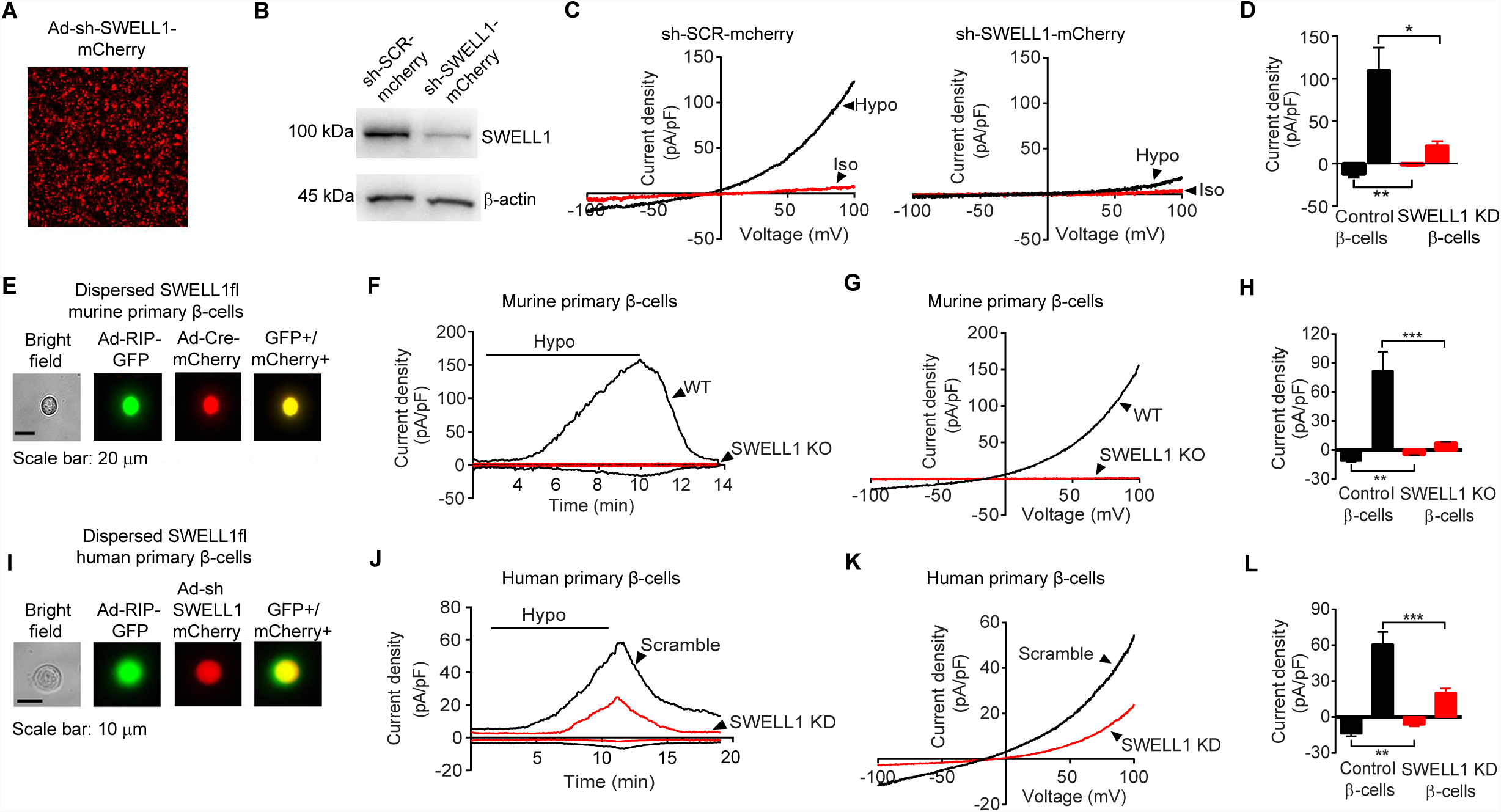
*SWELL1* mediates VRAC in MIN6 cells and primary mouse and human pancreatic β-cells. **a**, mCherry fluorescence of the mouse insulinoma cell line (MIN6) transduced with an adenovirus expressing a short-hairpin RNA directed to SWELL1 (shSWELL1-mCherry). **b**, SWELL1 Western blot in MIN6 cells transduced with shSWELL1 compared to scrambled short-hairpin RNA (shSCR). β-actin loading control (Supplementary Fig. 5 for full blots). **c**, Current-voltage relationship of VRAC in MIN6 cells at baseline (black) and after hypotonic swelling (Hypo, 210 mOsm, blue) after adenoviral transduction with shSCR (left) and shSWELL1 (right). **d**, Mean current inward and outward densities at +100 and -100 mV (n_shSCR =_ 3 cells; n_shSWELL1_ = 4 cells). **e**, Bright-field, GFP, mCherry and merged images of freshly dispersed primary β-cells from *SWELL1^fl/fl^* mouse islets co-transduced with Ad-RIP1-GFP and Ad-Cre-mCherry. (**f-g)**, Current-time relationship (**f**) and current-voltage relationship (**g**) of swell-activated VRAC in wild-type (WT: Ad-mcherry/SWELL*^fl/fl^*) and SWELL1 knockout (KO: Ad-Cre-mCherry/*SWELL1^fl/fl^*) mouse primary β-cells. (**h**) Mean current inward and outward densities at +100 and -100 mV (n = 5 cells, each group). (**i**) Bright-field, GFP, mCherry and merged images of freshly dispersed primary β-cells from human islets co-transduced with Ad-RIP1-GFP and Ad-shSWELL1-mCherry. (**j**) Current-time relationship and (**k**) current-voltage relationship of swell-activated VRAC in WT and SWELL1 knockdown primary human β-cells. (**l**) Mean current inward and outward densities at +100 and -100 mV (n_shSCR_ = 5 cells; n_shSWELL1_ = 10 cells). Ramp protocol is from +100 mV to -100 mV (ramp duration: 600 ms, holding potential: 0 mV). Error bars in d, h, and l denote ± s.e.m. * P < 0.05; **P < 0.01; *** P < 0.001, unpaired t-test.

Having established that SWELL1 is required for this previously enigmatic depolarizing swell-activated Cl^−^ current16,17 in MIN6 cells, and in both mouse and human primary β-cells we asked whether physiologic glucose-mediated β-cell swelling^18^ is sufficient to activate SWELL1-mediated VRAC. First, we measured β-cell size by light microscopy in WT and SWELL1-deficient primary murine and human β-cells in response to glucose-stimulated swelling (at 35-37°C). WT murine β-cells swell 6.8 ± 1.6 % in cross-sectional area upon perfusion of 16.7 mM glucose (from 1 mM basal glucose) and reach a maximum size at 12 minutes post glucose-stimulation, followed by a reduction in β-cell size (Fig. 2a), consistent with regulatory volume decrease (RVD). In contrast, SWELL1 KO murine β-cells swell monotonically to 8.2 ± 2.4 % and exhibit no RVD (Fig. 2a). WT human β-cells show a similar trend, swelling 8.6 ± 3.5 %, followed by RVD (Fig. 2b). SWELL1 KD human β-cells swell monotonically to 6.0 ± 1.5 % (Fig. 2b), and similar to SWELL1 KO murine β-cells (Fig. 2a), exhibit no RVD. These data indicate that physiological increases in glucose induce β-cell swelling and that SWELL1 is required for RVD in primary β-cells, as observed in cell lines^11,12,19^. Next, we applied the perforated-patch clamp technique to primary β-cells at 35-37°C in order to measure currents under the same physiological conditions that induce glucose-mediated β-cell swelling. We find that β-cell VRAC activates in response to physiological increases in glucose in MIN6 cells (Supplementary Fig. 2) and in both mouse (Fig. 2c-d) and human (Fig. 2e-f) β-cells, and is blocked by the selective VRAC inhibitor, DCPIB. Importantly, the time-course of VRAC activation in β-cells either tracks or lags the latency of β-cell swelling in response to stimulatory glucose (~8 minutes), consistent with a mechanism of glucose-mediated β-cell swell-activation of SWELL1-mediated VRAC. Thus, SWELL1-mediates a glucose-sensitive swell-activated depolarizing Cl^−^ current in β-cells.

**Figure 2.**
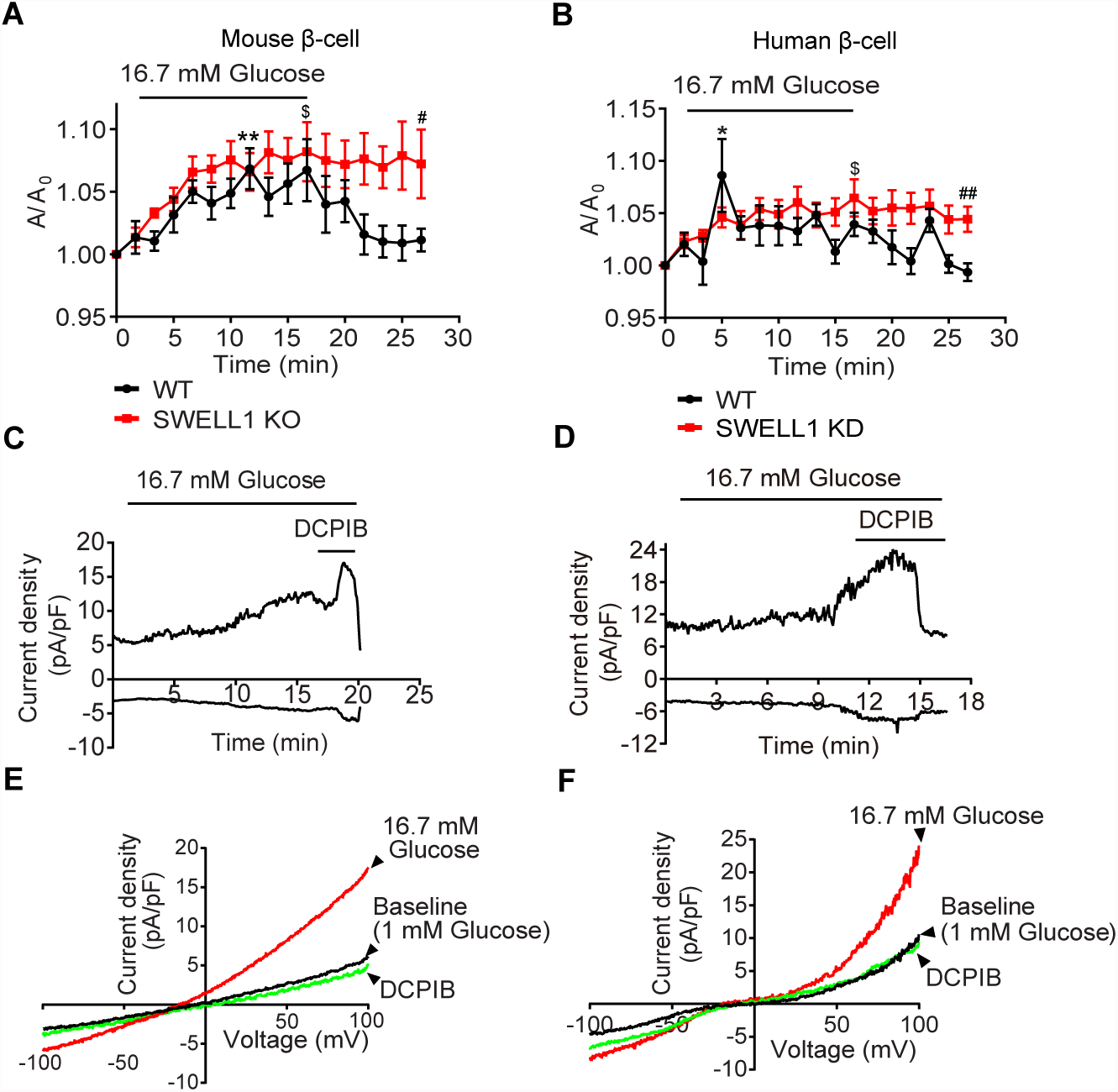
β-cell VRAC is activated by physiological swelling in response to glucose stimulation. **a-b**, Cross-sectional area of primary WT (n = 12) and SWELL1 KO (n = 7) murine β-cells (**a**) and WT (n = 9) and SWELL1 KD (n = 8) human β-cells (**b**) in response to glucose-stimulation (1, 16.7, 1 mM glucose). **c-d**, Murine primary β-cell VRAC over time (**c**) and current-voltage relationship (**d**) with DCPIB inhibition (10 uM). **e-f**, Human primary β-cell VRAC over time (**e**) and current-voltage relationship (**f**) with DCPIB inhibition (10 uM). Recordings in **c-f** performed at 35-37°C in perforated-patch configuration. In (a), **P < 0.01 vs 0 min in WT, paired t-test; ^$^P < 0.05 vs 0 min in SWELL1 KO, paired-test; ^#^P < 0.05 WT vs SWELL1 KO, unpaired t-test. In (b), *P < 0.05 vs 0 min in WT, paired t-test; ^$^P < 0.05 vs 0 min in SWELL1 KD, paired-test; ^##^P < 0.01 WT vs SWELL1 KD, unpaired t-test.

To determine whether SWELL1-mediated depolarizing Cl^−^ current is required to depolarize the β-cell membrane potential to the activation threshold of voltage-gated Ca^2+^ channels (VGCC) essential for insulin granule fusion, we next measured glucose-stimulated intracellular Ca^2+^ in WT and SWELL1 - deficient MIN6 β-cells, primary mouse and human β-cells. Using CRISPR/cas9 technology, we generated multiple SWELL1 KO MIN6 cell lines (Supplementary Fig. 3), confirming *SWELL1* gene disruption by PCR (Supplementary Fig. 3a), SWELL1 protein deletion (Fig. 3a & Supplementary Fig. 3b) and ablation of SWELL1-mediated current (Fig. 3b) in these cells. We find that glucose-stimulated Ca^2+^ transients are entirely abolished in SWELL1 KO MIN6 compared to WT cells (Fig. 3c-e), despite preserved KCl (40 mM) stimulated Ca^2+^ transients (control for intact β-cells excitability). Co-application of a selective VGCC blocker nifedipine (10 μM) fully inhibits these glucose-stimulated Ca^2+^ transients in WT MIN6 cells, consistent with a mechanism of membrane depolarization and VGCC activation (Supplementary Fig. 4a&b).

**Figure 3.**
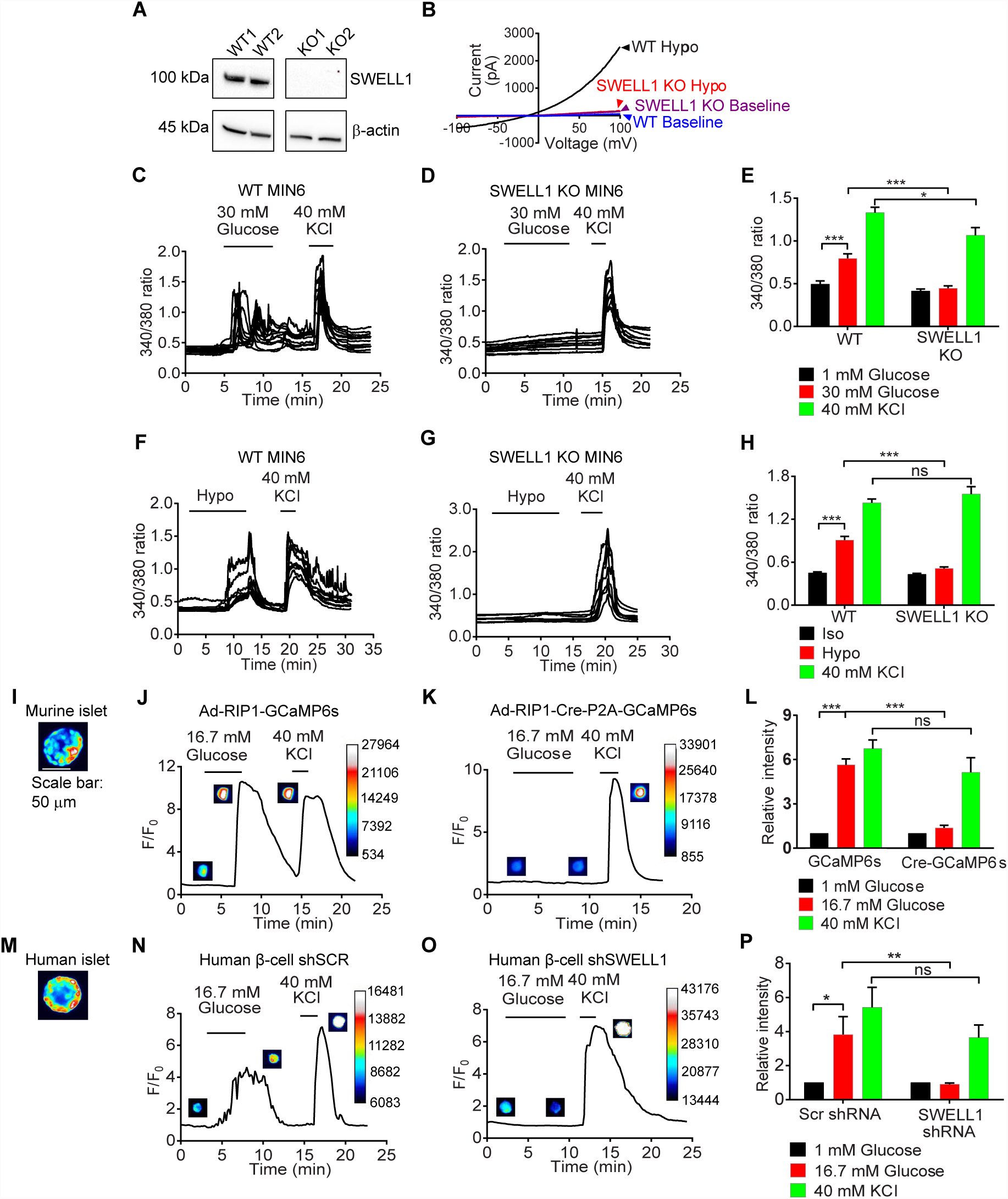
*SWELL1-*mediated VRAC is required for both glucose- and hypotonic swelling induced Ca^2+^ signaling in β-cells. **a**, SWELL1 Western blot in WT and CRISPR/Cas9-mediated SWELL1 KO MIN6 cell lines (Supplementary Fig. 5 for full blots). **b**, Current-voltage plots of SWELL1-mediated current in response to hypotonic swelling in WT and SWELL1 KO MIN6 cells confirm complete ablation of hypotonic swelling stimulated VRAC in SWELL1 KO MIN6 cells. (**c-d**) Fura-2 Ca^2+^ transients in WT (**c**) and SWELL1 KO (**d**) MIN6 cells in response to 30 mM glucose-stimulation (basal 1 mM glucose). 40 mM KCl stimulation confirms cell viability and excitability. (**e**) Mean peak Fura-2 ratio in glucose-stimulated WT (n = 34) and SWELL1 KO (n= 36) MIN6 cells. (**f-g**) Fura-2 Ca^2+^ transients in WT (**f**) and SWELL1 KO (**g**) MIN6 cells in response to swell-stimulation (210 mOsm; isotonic 300 mOsm). (**h**) Mean peak Fura-2 ratio in swell-stimulated WT (n = 42) and SWELL1 KO (n = 26) MIN6 cells. (**i**) Ad-RIP1-GCaMP6s transduced SWELL1^fl/fl^ murine islet. (**j-k**) GCaMP6s Ca^2+^ transients in WT (J, Ad-RIP1-GCaMP6s/SWELL1^fl/fl^) and SWELL1 KO (K, Ad-RIP1-Cre-P2A-GCaMP6s/SWELL1^fl/fl^) primary murine β-cell in response to 16.7 mM glucose-stimulation (basal 1 mM glucose). 40 mM KCl stimulation. Insets show β-cell fluorescence images at the indicated times in the experiment. (**l**) Mean peak values of GCaMP6s Ca^2+^ transients from Ad-RIP1-GCaMP6s/SWELL1^fl/fl^ (n = 14) and Ad-RIP1-Cre-P2A-GCaMP6s/SWELL1^fl/fl^ (n = 10). (**m**) Ad-RIP1-GCaMP6s transduced human islet. (**o-p**) GCaMP6s Ca^2+^ transients in human primary β-cell co-transduced with Ad-RIP1-GCaMP6s+Ad-shSCR-mCherry (**o**) and Ad-RIP1-GCaMP6s+Ad-shSWELL1-mCherry (**p**) in response to 16.7 mM glucose-stimulation (basal 1 mM glucose). 40 mM KCl stimulation. (**q**) Mean peak values of GCaMP6s Ca^2+^ transients from shSCR (n = 6) and shSWELL1 (n = 8). Error bars represent mean ± SEM, **p < 0.01, ***p < 0.001.

As β-cells are also known to depolarize, fire Ca^2+^ transients, and secrete insulin via a glucose-independent hypotonic swelling mechanism^16,20^, we next examined swell-induced Ca^2+^-signaling in β-cells in response to hypotonic stimulation (220 mOsm) in the absence of glucose-stimulation (0 mM glucose). We find that hypotonic swelling alone can trigger robust Ca^2+^ transients in WT MIN6 cells (Fig. 3f&h) and these elevations in cytosolic Ca^2+^ recover rapidly upon restoration of isotonic solution (Fig. 3f). In contrast, SWELL1 KO MIN6 cells are entirely non-responsive to hypotonic swelling induced Ca^2+^ transients (Fig. 3f-h), despite preserved KCl stimulated Ca^2+^ responses, consistent with SWELL1-mediating a glucose-independent, swell-activated depolarizing current in β-cells. Similar to the case with glucose-stimulated Ca^2+^ signaling, we also find that hypotonic swelling triggered Ca^2+^ transients are fully inhibited by VGCC blockade (Supplementary Fig. 4c&d), implicating β-cell membrane depolarization followed by VGCC activation, as opposed to alternative hypo-osmotically activated Ca^2+^ influx pathways (i.e. TRP channels)^21^.

**Figure 4.**
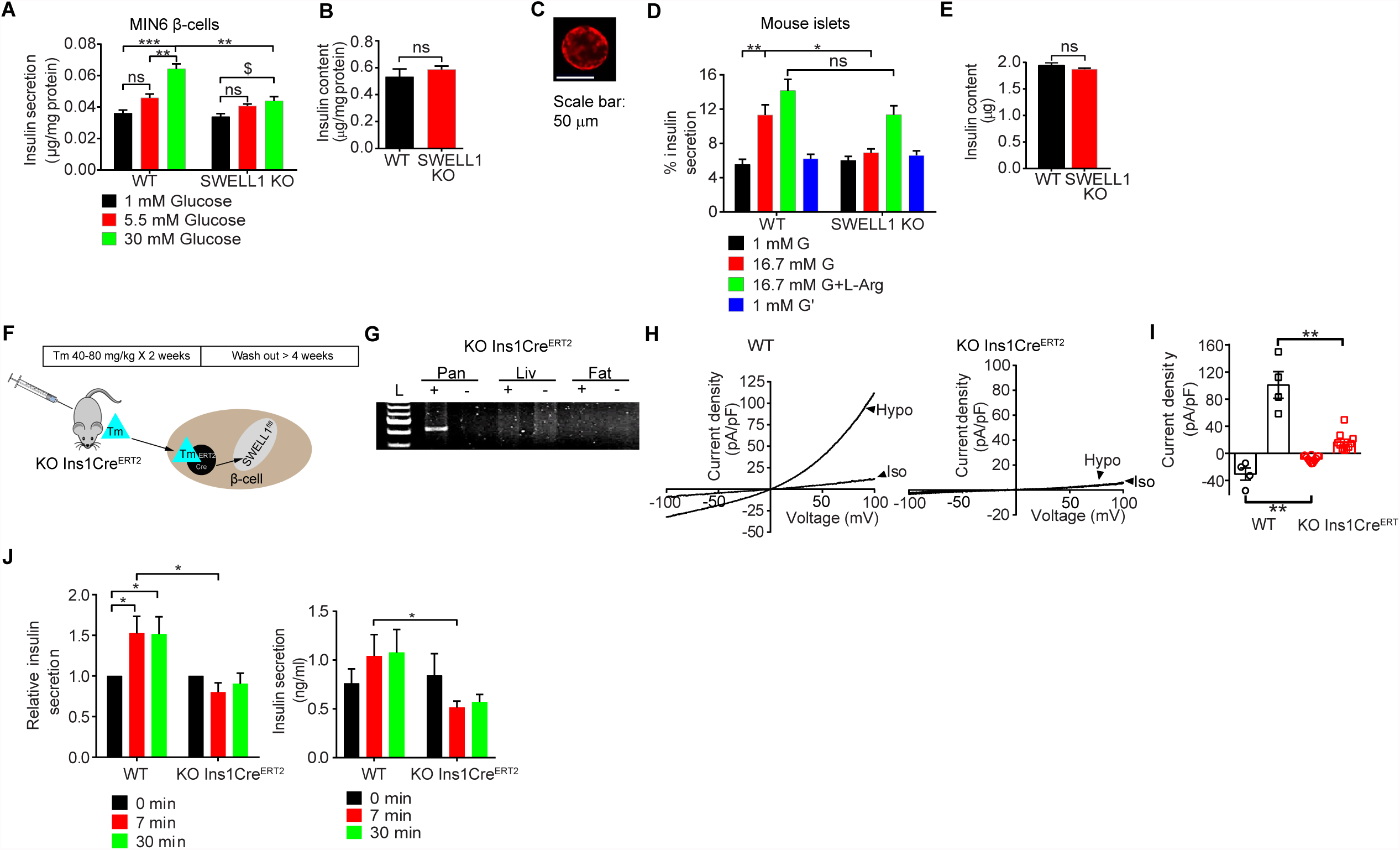
β-cell *SWELL1* is required for glucose-stimulated insulin secretion *in vitro* and *in vivo*. **a**, Glucose-stimulated insulin secretion (GSIS) in WT and SWELL1 KO MIN6 cells in response to 1, 5.5 and 30 mM glucose (n = 3, each). **b**, Total insulin content normalized to total MIN6 cell protein content (n = 3, each). **c**, Representative image of Ad-RIP1-Cre-RFP transduced mouse islet. **d**, GSIS in WT (Ad-RIP1-RFP/*SWELL1^fl/fl^*) and β-cell targeted SWELL1 KO (Ad-RIP1-Cre-RFP/*SWELL1^fl/fl^*) islets. Insulin secretion is expressed as percentage of the total insulin content (n = 4, each). **e**,Total insulin content of WT and β-cell targeted SWELL1 KO islets (n = 4, each). (**f**) Tamoxifen-inducible β-cell-targeted targeted *SWELL1* inactivation using *Ins1Cre^ERT2^* mouse crossed with *SWELL1^fl/fl^* mice (*KOIns1Cre^ERT2^*). (**g**) PCR across *SWELL1* Exon 3 in genomic DNA extracted from pancreas, liver and adipose tissues of a *KO Ins1Cre^ERT2^* mouse treated with (+) and without (-) tamoxifen. *Cre*-mediated *SWELL1* recombination results in a 426 bp amplicon. (**h**) Current-voltage relationship of swell-activated VRAC in mouse primary β-cells isolated from *SWELL1^fl/fl^* +tamoxifen (WT) and *KO Ins1Cre^ERT2^* +tamoxifen (*KO Ins1Cre^ERT2^*). (**i**) Mean inward and outward current densities at +100 and -100 mV (n = 4 cells for WT and n = 10 cells for *KO Ins1Cre^ERT2^*). (**j**) Glucose-stimulated insulin secretion in WT and *KO Ins1Cre^ERT2^* mice in response to 2g/kg glucose in the first 30 min (n = 8, male, for each group). Error bars represent mean ± SEM, * p < 0.05; **p < 0.01; ***p < 0.001 vs. WT; ns: not statistically significant, one-way ANOVA for in-group comparison, unpaired t-test for between-group comparison. For (**j**), non-parametric one-way ANOVA (Kruskal Wallis test) was performed. Error bars represent mean ± SEM, * p < 0.05; **p < 0.01; ***p < 0.001 vs. WT (30 mM glucose); ns: not statistically significant

We next examined SWELL1-dependent Ca^2+^ signaling in primary mouse and human β-cells. We generated adenoviruses expressing the genetically encoded Ca^2+^-sensor GCaMP6s under control of the rat insulin promoter 1 (RIP1), either alone (Ad-RIP1-GCaMP6s), or in combination with Cre-recombinase (Ad-RIP1-Cre-P2A-GCaMP6s). This approach provides a robust β-cell restricted fluorescent Ca^2+^ sensor while simultaneously allowing for β-cell targeted Cre-mediated *SWELL1* deletion in cultured *SWELL1^fl/fl^* islets isolated from *SWELL1^fl/fl^* mice (Fig. 3i). GCaMP6s Ca^2+^ imaging reveals robust glucose-stimulated Ca^2+^ transients in freshly dissociated WT primary murine β-cells (Ad-RIP1-GCaMP6s/*SWELL1^fl/fl^*; Fig. 3j&l) and these are significantly suppressed in SWELL1 KO β-cells (Ad-RIP1-Cre-P2A-GCaMP6s/*SWELL1^fl/fl^*; Fig. 3k&l), despite preserved KCl stimulated Ca^2+^ responses. We used a similar approach in human islets, whereby we co-transduced islets with Ad-RIP1-GCaMP6s (Fig. 3m) and either Ad-U6-shSWELL1-mCherry or Ad-U6-shSCR-mCherry. Upon islet dissociation, we imaged only double-labelled GCaMP6s+/mCherry+ primary human β-cells. As with mouse primary β-cells, we observe robust glucose-stimulated Ca^2+^ transients in Ad-shSCR treated human primary β-cells (Fig. 3n&p) and this is markedly abograted upon Ad-shSWELL1-mediated SWELL1 knock-down (Fig. 3o&p). Collectively, these data demonstrate that SWELL1 is required for both glucose- and swell-activated Ca^2+^ signaling in MIN6 cells and in mouse and human primary β-cells. Moreover, these data suggest that the depolarizing SWELL1-mediated Cl^−^ current is necessary for β-cell depolarization in response to glucose-stimulation. In pancreatic β-cells, physiological intracellular Cl^−^ concentration is maintained at 34-36 mM^22,23^ by NKCC1 transporters^24,25^ to generate a depolarizing Cl^−^ current upon activation of a Cl^−^ conductance, since E_Cl-_ = ~-35 mV. Therefore, NKCC1 blockade by bumetanide^26,27^ is predicted to reduce intracellular Cl^−^, drop E_cl-_ and thereby diminish or abolish β-cell membrane depolarization by a SWELL1-mediated glucose-stimulated Cl^−^ conductance. Consistent with this prediction, we find that application of bumetanide (10 μM) fully inhibits glucose-stimulated Ca^2+^ signaling in both WT MIN6 cells (Supplementary Fig. 4e&f) and WT primary murine β-cells (Supplementary Fig. 4g&h).

To determine the impact of SWELL1-dependent glucose-stimulated Ca^2+^ signaling on insulin secretion in β-cells we measured glucose-stimulated insulin secretion (GSIS) in WT and SWELL1 KO MIN6 cells. We find that the glucose-dependent increase in insulin secretion in WT MIN6 cells is significantly diminished in SWELL1 KO MIN6 cells (Fig. 4a), particularly at higher glucose concentration (30 mM), despite no change in total insulin content ( Fig. 4b). We next isolated islets from *SWELL1^fl/fl^* mice followed by transduction with either Ad-RIP1-RFP (WT) or Ad-RIP1-Cre-P2A-RFP (SWELL1 KO; Fig. 4c). Similar to MIN6 cells, we note a significant reduction in GSIS (16.7 mM glucose) in SWELL1 KO compared to WT islets (Fig. 4d), despite relatively preserved L-arginine stimulated insulin secretion (Fig. 4d), and similar total insulin content (Fig. 4e).

We next generated tamoxifen-inducible β-cell-targeted *SWELL1* KO mice by crossing *SWELL1^fl/fl^* mice with *Ins1Cre^ERT2^* mice (*KO Ins1Cre^ERT2^*, Fig. 4f)^28^ to examine the requirement of β-cell SWELL1 for insulin secretion *in vivo*. After tamoxifen-administration (40-80 mg/kg/day x 5 days), we observe pancreas-restricted *SWELL1* recombination in *KO Ins1Cre^ERT2^* mice by PCR across *SWELL1* Exon 3 (426 bp amplicon; Fig. 4g) and complete ablation of SWELL1-mediated current in 80% of patch-clamped β-cells (8/10 cells; Fig. 4h-i), while 100% of β-cells from tamoxifen-induced *SWELL1^fl/fl^* β-cells (WT) had robust hypotonically-activated SWELL1-mediated currents (4/4 cells). We find that basal (fasting) serum insulin are similar between WT and *KO Ins1Cre^ERT2^* mice (Fig. 4j), which is consistent with preserved basal insulin secretion observed at low glucose *in vitro* in SWELL1 KO MIN6 cells (Fig. 4a) and in β-cell-targeted SWELL1 KO murine islets (Fig. 4d). However, with glucose-stimulation (2 g/kg i.p.) we find that insulin secretion is significantly impaired in *KO Ins1Cre^ERT2^* mice compared to WT mice (Fig. 4j). Overall, these data are consistent with a requirement of β-cell SWELL1 for glucose-stimulated, Ca^2+^-dependent insulin secretion both *in vitro* and *in vivo*.

VRAC/I^Cl,SWELL^ has been studied for decades through electrophysiological recordings in numerous cell types^29^^−^^31^, but only recently has it been discovered that SWELL1/LRRC8a, and associated LRRC8 isoforms b-e, form the VRAC channel complex in common cell lines^11-13.^ Accordingly, the physiological role of SWELL1-mediated VRAC in primary cells remains unexplored. We recently showed that SWELL1 is required for VRAC in adipocytes where it senses adipocyte hypertrophy in the setting of obesity and regulates insulin-PI3K-AKT2-GLUT4 mediated glucose uptake and systemic glycemia ^32^. Here we asked whether SWELL1 is required for VRAC described previously in the pancreatic β-cell^16^-^18^ and whether the VRAC hypothesis ^33^ can be explained by a putative glucose-mediated swell sensing function of the SWELL1/LRRC8 channel complex in β-cells. Indeed, our data are consistent with a model in which SWELL1 is a required component of a swell-activated depolarizing Cl^−^ channel that activates in response to glucose-stimulated β-cell swelling and is required for membrane depolarization, VGCC activation, Ca^2+^-mediated insulin vesicle fusion and insulin secretion. In this model, the hyperpolarizing K^+^ conductances ^4^^−^^10^ act as a “brakes” on β-cell excitability and insulin secretion, while SWELL1-mediated VRAC is the “accelerator” -promoting β-cell excitability in response to glucose-mediated β-cell swelling. Overall, these data suggest that β-cell SWELL1 acts as a glucose sensor by coupling β-cell swelling to β-cell depolarization - a form of swell-activation or swell-secretion coupling - to potentiate glucose-stimulated insulin secretion. In the broader context of our findings on SWELL1 signaling in the adipocyte ^32^, these data suggest that SWELL1 coordinately regulates both insulin secretion and insulin sensitivity ^32^ in response to a nutrient load, highlighting the importance SWELL1 in the regulation of systemic glucose metabolism.

## AUTHOR CONTRIBUTIONS

Conceptualization, R.S.; Methodology, C.K., S.B.S., A.W.N., A.M., Y.Z., L.X., S.K.G., S.P. Y.G. R.S.; Formal Analysis, C.K., R.S.; Investigation, C.K., S.S., A.M., S.P., Y.G.; Resources, R.S., A.W.N.; Writing – Original Draft, R.S., Writing – Review & Editing, R.S., C.K., S.B.S, A.W.N.; Visualization, C.K., R.S.; Supervision, R.S.; Funding Acquisition, R.S.

## ACKNOWLEDGMENTS

We thank Dr. John Engelhardt for sharing human islets obtained from the Integrated Islet Distribution Program (IIDP) and Dr. Yumi Imai sharing human islets obtained from Prodo Laboratories. We thank Shanming Hu for assisting with mouse islet isolations and Dr. Robert Tsushima for thoughtful reading of the manuscript and comments. This work was supported by grants from the NIH NIDDK 1R01DK106009 (R.S.), R01DK097820 (A.W.N.), R24DK96518 (A.W.N.) and the Roy J. Carver Trust (R.S.).

---

**Supplementary Figure 1: Fluorescence images of adenovirally transduced murine and human islets**

**a**, Murine islets freshly isolated from SWELL1^fl/fl^ mice, cultured (BF: Bright field) and then co-transduced with Ad-RIP1-GFP (GFP) and Ad-CMV-mCherry (top, cytosolic mCherry: Control) or Ad-CMV-Cre-mCherry (bottom: nuclear-localized Cre-mCherry fusion protein; SWELL1 KO). **b**, Human islets cultured (BF: Bright field) and then co-transduced with Ad-RIP1-GFP (GFP) and Ad-U6-shSCR-mCherry (top, mCherry: Control) or Ad-U6-shSWELL1-mCherry (bottom: mCherry; SWELL1 KD). Scale bar represents 50 μm.

**Supplementary Figure 2: MIN6 β-cell VRAC is activated by glucose stimulation**

Glucose-stimulated VRAC recorded in patch-clamped MIN6 β-cell upon application of 30 mM glucose from a basal glucose of 1 mM. Recordings performed at 35-37°C in perforated-patch configuration.

**Supplementary Figure 3: CRISPR/cas9-mediated *SWELL1* ablation in MIN6 cells**

**a**, Guide RNA sequences targeting exon 3 of the *SWELL1* gene were used in combinations of either 1A+2B or 2B+3C to generate KO1 and KO2 clones respectively. Upon interacting with *cas9* enzyme and corresponding guide pairs the target region undergoes non-homologous end joining (NHEJ) repair. This results in the deletion of DNA base pairs in-between the two target guide sites. Using specific primers for the regions flanking the two target guide sites, the wildtype, WT (non-transfected) cells generate a fragment of size 661 and 879 bps for the 1A/2B and 2B/3C sites respectively, upon PCR amplification. The KO1 and KO2 clones (transfected) generate a deleted DNA fragment of size approximately 410 and 639 bps for the 1A/2B and 2B/3C sites respectively. In the agarose gel image, the DNA fragment sizes are indicated in base-pairs (bp) and ‘L’ indicates ladder. **b,** Western-blot of SWELL1 protein in WT (non-transfected, WT1, WT2) and multiple SWELL1 KO MIN6 clones (KO1-5). *β*-actin is used as the loading control (Supplementary Fig. 5 for full blots).

**Supplementary Figure 4: Glucose and hypotonic swell-stimulated Ca^2+^ signaling is dependent on VGCC and elevated intracellular chloride in β-cells**

**a**, Fura-2 Ca^2+^ transients in WT MIN6 cells in response to 30 mM glucose-stimulation (basal 1 mM glucose) in the presence of VGCC blocker nifedipine (10 μM). **b**, Mean peak Fura-2 ratio from **a** (n = 16 cells). **c**, Fura-2 Ca^2+^ transients in WT MIN6 cells in response to hypotonic swelling (210 mOsm; isotonic 300 mOsm) in the presence of VGCC blocker nifedipine (10 μM). **d**, Mean peak Fura-2 ratio from **c** (n = 43 cells). **e**, Fura-2 Ca^2+^ transients in WT MIN6 cells in response to 30 mM glucose-stimulation (basal 1 mM glucose) in the presence of NKCC1 inhibitor, bumetanide (10 μM). **f**, Mean peak Fura-2 ratio from **e** (n = 46 cells). **g**, GCaMP6s Ca^2+^ transients in WT (Ad-RIP1-GCaMP6s/SWELL1^fl/fl^) primary murine β-cell in response to 16.7 mM glucose-stimulation (basal 1 mM glucose) in the presence of NKCC1 inhibitor, bumetanide (10 μM). **h**, Mean peak values of GCaMP6s Ca^2+^ transients from **g** (n = 7 cells). In all experiments, 40 mM KCl stimulation confirms cell viability and excitability. Error bars represent mean ± s.e.m., **p < 0.01; ***p < 0.001; ns: not statistically significant.

